# Prioritizing Parkinson’s Disease genes using population-scale transcriptomic data

**DOI:** 10.1101/231001

**Authors:** Yang I Li, Garrett Wong, Jack Humphrey, Towfique Raj

**Author notes:** These authors contributed equally to this work (Yang I Li, Garrett Wong). Correspondence should be addressed to T.R.

## Abstract

Genome-wide association studies (GWAS) have identified over *41* susceptibility loci associated with late-onset Parkinson’s Disease (PD) but identifying putative causal genes and the underlying mechanisms remains challenging. To address this, we leveraged large-scale transcriptomic datasets to prioritize genes that are likely to affect PD. We found *29* gene associations in peripheral monocytes, and *44* gene associations whose expression or differential splicing in prefrontal cortex is associated with PD. This includes many novel genes but also known associations such as MAPT, for which we found that variation in exon *3* splicing explains the common genetic association. Genes identified in our analyses are more likely to interact physically with known PD genes and belong to the same or related pathways including lysosomal and innate immune function. Overall, our study provides a strong foundation for further mechanistic studies that will elucidate the molecular drivers of PD.

## Introduction

Parkinson’s disease (PD) is the second most common neurodegenerative disorder after Alzheimer’s disease (AD). PD is characterized by the formation of intracytoplasmic inclusions known as Lewy bodies containing α-synuclein (α-syn) and by the loss of dopaminergic neurons primarily in the substantia nigra^1,2^. Over the last decade a number of genetic susceptibility factors have been identified for PD including nine genes linked to heritable, monogenic forms of PD ^2^. More recently, over 41 genetic susceptibility loci have been associated with late-onset PD in the largest GWAS meta-analysis of PD to date^3^. Within these risk loci, a few genes have been identified as potentially causal, but for the majority of loci, it is not yet known which genes underlie PD risk. More generally, it is currently unclear whether PD risk genes identified thus far by GWASs belong to coherent pathways such as those involved in lysosomal and autophagy function ^3^ or whether PD risk genes belong to broader pathways including innate and adaptive immune function.

The integration of large-scale functional genomic data with PD GWAS has been a powerful approach to characterize the functional effects of PD-associated variants^4,3^ For example, we have previously shown that PD-associated loci are enriched in expression quantitative loci (eQTLs) from peripheral monocytes but not CD4^+^ T-cells ^5^. Similarly, recent studies used cell-type specific functional annotations to show that genes in PD GWAS loci are often preferentially expressed in CD14+ monocytes^6,7^. These findings support the involvement of the innate immune cells (peripheral monocytes, central nervous system macrophages and/or microglia) in the etiology of PD.

Many studies have also attempted to use functional genomics data in postmortem human brains to ascribe function to PD risk variants^8,9,10^. Generally, these were underpowered due to limited sample sizes and the high cellular heterogeneity of brain tissues, but in a few cases they have led to novel biological hypotheses about the mechanisms underlying PD-associated genetic associations. For instance, in the *MAPT* region, the absence of a well-characterized inversion at 17q21 marks the PD-associated H1 haplotype, which has been reported to be associated with exon 10 inclusion^11^, *MAPT* exon 3 exclusion^12^, and also increased total *MAPT* expression^11^. While these conflicting reports suggest that *MAPT* is involved in PD, it is unknown what specific mechanism drives the association to PD and whether the H1 or H2 haplotypes are also associated with differential expression of other genes in the locus.

Here, we leveraged large-scale transcriptomics data to identify candidate PD genes and to better understand the primary mechanisms that underlie PD genetic risk factors. We first used broad atlases of gene expression that include brain^13,14^ and immune cell-types^15,14^ to find tissues with specifically expressed genes enriched PD susceptibility loci. We then leveraged large-scale transcriptomic data from primary monocytes^5,16,17^ and a large-scale dorsolateral prefrontal cortex dataset^18^ to perform transcriptome-wide association study of PD. We prioritized 66 genes, whose predicted expression or splicing levels in peripheral monocytes cells and in DLPFC are significantly associated with PD risk. Overall, our results support the importance of innate immune and lysosomal-related pathways in PD etiology, and suggest specific genes and mechanisms at each GWAS locus as determinants of PD risk.

## Results

### Heritability enrichment of expressed genes identifies PD-relevant tissues

To obtain a better understanding of how genetic variants affect PD risk, we first attempted to identify tissues likely to be relevant in PD pathology. To this end, we used LD score regression for specifically expressed genes (LDSC-SEG), which identifies tissues in which genes with increased expression are enriched in SNPs that tag a large amount of PD heritability^19^. When applied to 53 tissues from the GTEx Consortium^13^, we detected an enrichment at a 5% FDR threshold (— log_10_ p-value > 2.86) for six tissues including the amygdala, substantia nigra, anterior cingulate cortex, frontal cortex, hypothalamus, and cervical (C1) spinal cord (Fig. 1b). These findings can be replicated using a larger expression panel comprising of 152 cell-types^14^ (Supplementary Fig. 1) and suggest that central nervous system (CNS) tissues are major players in PD. In contrast, there is no enrichment of Alzheimer’s disease (AD) SNP heritability near genes expressed in these CNS tissues. This may suggest fundamentally different roles for the various neuronal cell-types in the two neurodegenerative diseases. When we used LDSC-SEG on expression data from primary mouse CNS cells^20^, we found that neurons and oligodendrocytes, but not astrocytes, preferentially expressed genes enriched in SNP heritability for PD pathology (Supplementary Fig. 2). Finally, we applied LDSC-SEG on an atlas of 291 mouse immunological cell types^15^ in order to verify whether PD signal enrichment near genes specifically expressed in myeloid cells are higher than compared to other immune cell types, as previous results in the field suggest^7,6^. However, unlike in AD, where there is a marked enrichment for myeloid cell types, there is a very modest enrichment in PD (Fig. 1b). It is possible that this lack of enrichment reflects a deficiency of annotating the genome by total expression tissue-specificity rather than, for example, the presence of mRNA isoforms or epigenetic marks.

**Figure 1:**
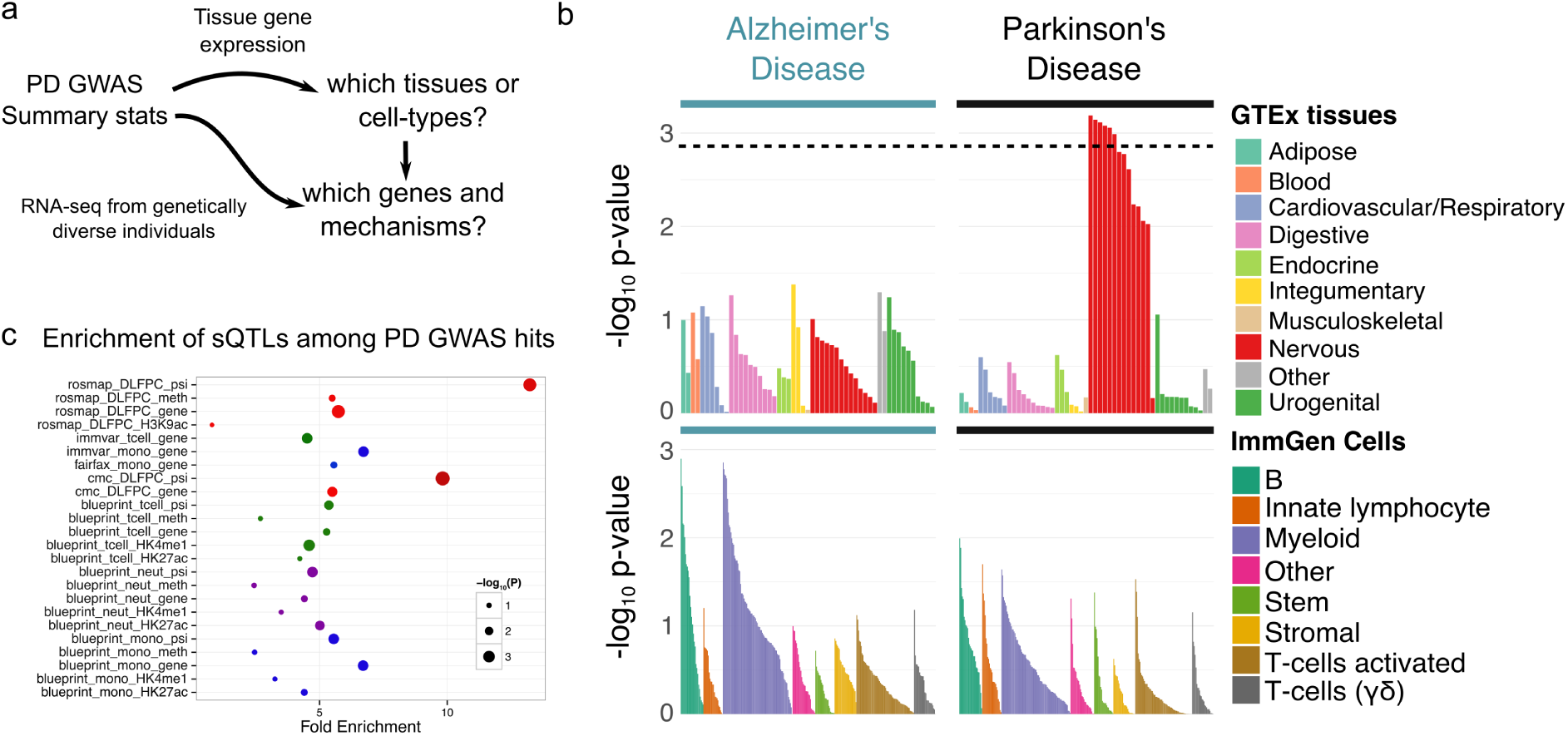
**(a)** Study rationale. To understand the genetic basis of PD, we leverage GWAS and large-scale gene expression datasets to: (1) identify tissues and cell-types enriched in GWAS signal, and (2) to identify genes and mechanisms through which PD associated loci *act* to affect disease risk. **(b)** LD-SEG analysis of AD and PD GWAS. Regions of the genome with specific expression in nervous tissues are most enriched for PD GWAS signal, among 53 GTEx tissues. Among sorted immune cells from ImmGen, we observed an enrichment of B-cells, innate lymphocytes, and myeloid cells. **(c)** Gene expression, splicing, methylation and chromatin QTL enrichments in PD GWAS (P-value < 10^−5^) across tissues and cell-types.

We next asked whether PD-associated variants were enriched among molecular (including expression, splicing, histone, and DNA methylation) quantitative trait loci (QTLs) in dorsolateral prefrontal cortex (DLFPC)^21,22^and immune cell-types^5,23^ (Fig. 1c). Indeed, as expected^24^, we found that most molecular QTLs are enriched in PD-associated variants, which implies a regulatory impact for PD-associated variants. More interestingly, however, we found that genetic variants that affect RNA splicing were most enriched (14× fold-enrichment) in variants associated to PD at p-value < 10^−5^. This observation is consistent across two different DLFPC datasets^18,21^, and motivated us to explore the hypothesis that PD-associated variants may affect RNA expression or splicing of nearby genes in myeloid cells and in cells from the CNS.

### Transcriptome-wide Association Study of PD in Peripheral Monocytes

To identify genes that are associated with PD in immune cell types, we first performed a transcriptome-wide association study (TWAS)^25,26^using summary-level data from a large PD GWAS ^27^ and transcriptome panels from peripheral monocytes. Briefly, the TWAS approach uses information from the gene expression measured in a reference panel and the PD GWAS summary statistics to evaluate the association between the genetic component of expression and PD status (See Online Methods). More specifically, we built TWAS models using expression data from primary peripheral monocytes in three independent cohorts^5,16,17^. We found 29 genes with expression in monocytes significantly associated with PD, including several genes in novel PD loci but previously associated with AD, e.g. *CD33* and *PILRB*^28,29,30^. Interestingly, *CD33* protein levels have previously been shown to be affected by PD risk variants in monocytes^31^, and *PILRB* is a binding partner for *TYROBP* (*DAP12*), the main regulator of the immune/microglia network activated in AD^32^. This overlap supports the existence of shared genetic risk factors – that likely involve immune-mediated mechanisms – for the two neurodegenerative diseases^33,34^.

As expected, we also see significant associations with several genes previously known to play a role in PD, e.g. *SNCA* and *LRRK2* (Fig. 2a, Supplementary Table 1), suggesting that our predictions are reliable for many loci. The effects of a large fraction of variants are expected to be shared across tissues and cell-types ^35^, we therefore asked whether some risk loci show evidence of cell-type-specific effects in monocytes, but not in cells from the CNS. Indeed, we found that the PD risk allele at the *LRRK2* locus is associated with increased expression of *LRRK2* in monocytes (rs76904798, *p* = 2.93 × 10^−8^), but not in DLFPC (*p* = 0.98) (Fig. 2b). At this locus, the GWAS signal colocalizes with the nearby eQTL signal in monocytes with posterior probability 0.99 (Supplementary Fig. 3). Recent studies have shown than both *SNCA* and *LRRK2* are highly expressed in human microglia^34^, and that the expression levels of these two genes are elevated in peripheral monocytes of PD patients compared to age-matched controls^36,37^. We propose that peripheral monocytes play an important role in the progression of PD and/or may serve as a proxy for microglial activities within the brain.

**Figure 2:**
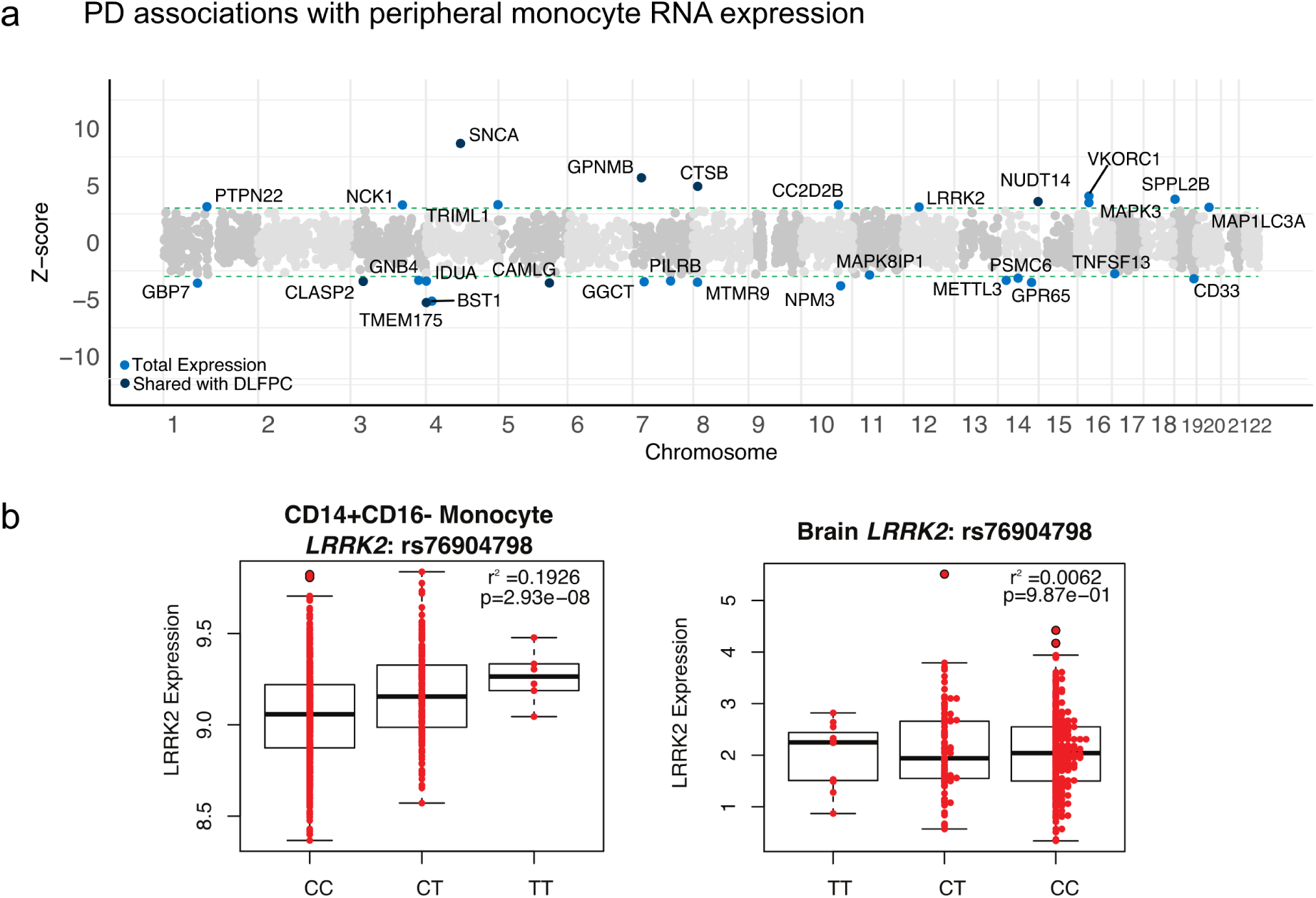
Contrasting contribution of peripheral monocytes and DLFPC to PD etiology. **(a)** TWAS of PD using gene expression from peripheral monocytes. **(b)** A SNP (rs76904798) that tags the PD GWAS risk loci at *LRRK2* is an eQTL that affects expression level of *LRRK2* in monocytes, but not in DLFPC.

### Transcriptome-wide Association Study of PD in DLFPC

Next, to identify candidate PD genes active in neuronal tissues, we used RNA sequencing data from DLPFC tissue from the CommonMind Consortium^18^, the largest publicly available brain tissue dataset of this kind, as a TWAS reference panel. In addition, because we were specifically interested in the role of RNA splicing in PD, we computed the “percent spliced in” (PSI) of splicing events that were detected in DLPFC RNA-seq using LeafCutter^38^, a method to detect and quantify alternative splicing events without a reference annotation. We used the PSI values for each splicing event to build our splicing TWAS model in addition to using total expression only, as commonly done in TWAS.

Our TWAS models for expression and splicing in the DLPFC identified several shared associations with our TWAS analysis in monocytes, but many are unique to either the monocytes or the DLPFC. In total, we found 18 and 26 genes that were associated to PD at 5% FDR in DLPFC through RNA expression and splicing, respectively. When we used coloc^39^ to assert colocalization between the PD association signal at these TWAS loci and expression or splicing QTLs, we found that a large number of loci (12 of 18, or 67%) had strong evidence of colocalization with DLPFC eQTLs, but only 4 of 26 (15%) for DLPFC sQTLs (Supplementary Table 1). The significantly lower colocalization between GWAS signals and sQTLs compared to eQTLs is unexpected, as we found higher enrichments for PD GWAS signal among sQTLs than eQTLs (Fig. 3c). A plausible explanation is that our TWAS model for RNA splicing produced a higher false positive rate than our TWAS model for RNA expression. However, another possibility is that genetic variants that affect RNA splicing tend to have smaller effects on complex traits or to be secondary associations. Approaches such as coloc depend on the alignment of strong effects both at the GWAS and QTL mapping levels, and thus they would be unable to detect colocalization between weak GWAS and/or sQTL signals. This is consistent with previous observations that eQTLs show stronger enrichments than sQTLs for top GWAS signals, but that sQTLs show stronger enrichments for variants with smaller but nonzero effects^24^. Indeed, when we plotted the posterior probabilities of our coloc analyses, we found that coloc performed well for eQTLs (Fig. 3b), assigning all probability density to H3 (independent signals) or H4 (colocalized signal). However, for a large number of sQTLs, coloc was underpowered to find evidence for colocalization (Fig. 3c), because it did not find evidence supporting the GWAS loci (H1), the sQTL association (H2), or neither (H0).

**Figure 3:**
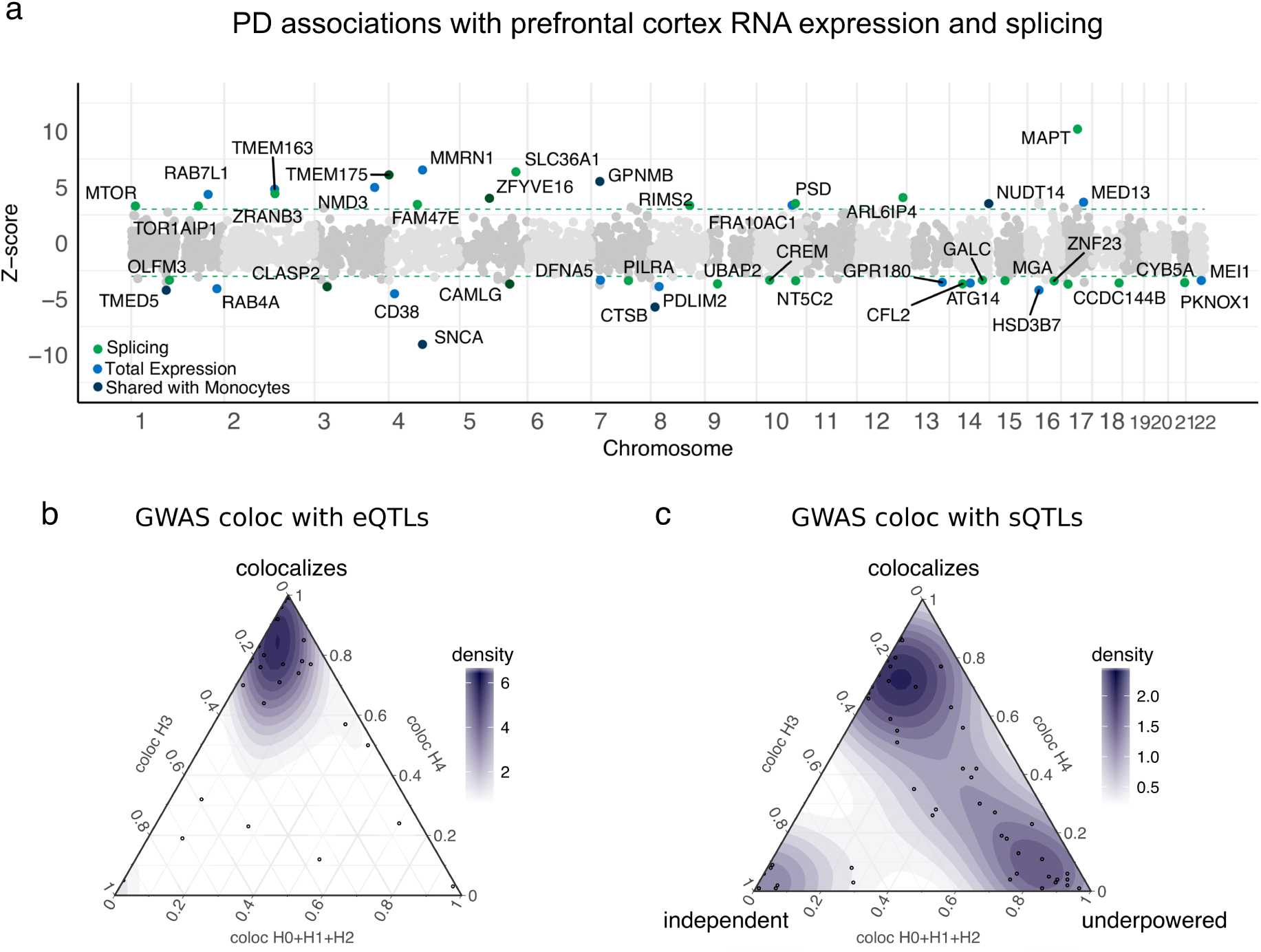
**(a)** PD TWAS using RNA expression and splicing measurements from post-mortem prefrontal cortices (CMC)^18^. Multiple association signals are shared between myeloid cells and prefrontal cortex, but differences suggest complementary contributions of both cell types. RNA splicing variation was also quantified using RNA-seq from prefrontal cortex samples. When genes are associated through both variation in expression and splicing, we used conditional analyses to identify which mechanism was most likely (Supplementary Materials). This revealed that the PD MAPT risk loci could be entirely explained by an effect of RNA splicing. **(b)** and **(c)** Ternary plots showing **coloc** posterior probabilities that TWAS loci found using RNA expression and splicing, respectively, belong to different sharing configurations. We considered H0+H1+H2 as evidence for lack of test power. H0: no causal variant, H1: causal variant for PD GWAS only, H2: causal variant for QTL only, H3: two distinct causal variants, H4: one common causal variant.

To further support the results from our splicing TWAS analysis, we performed additional analyses to replicate our TWAS in independent transcriptome ^21^ and GWAS datasets (23andMe cohort^27^). We first assessed whether the DLPFC TWAS results could be replicated in an independent transcriptomic reference panel by performing TWAS using DLPFC RNA-seq data from ROS/MAP ^22^. Twenty one of the 44 genes replicated at nominal *p* < 0.05 in the ROS/MAP TWAS. The direction of effect for most of the associations were concordant (Supplementary Table 2; Supplementary Fig. 4). Thus, the genetic effects on splicing are robust and unlikely to be due to artifacts unique to one of the datasets. To further replicate our findings, we repeated our TWAS using the CMC data and the PD GWAS summary statistics^27^ (using the 23andMe cohort only). When we compared the Z-scores obtained from these two TWASs, we found that 12 genes discovered in IPDGC (e.g. *MAPT*, *SNCA*, *ATG14*, *DVL3*, and *MTOR*) could be replicated (Fig. 4a). These two complementary replication efforts demonstrate the robustness of our TWAS results.

**Figure 4:**
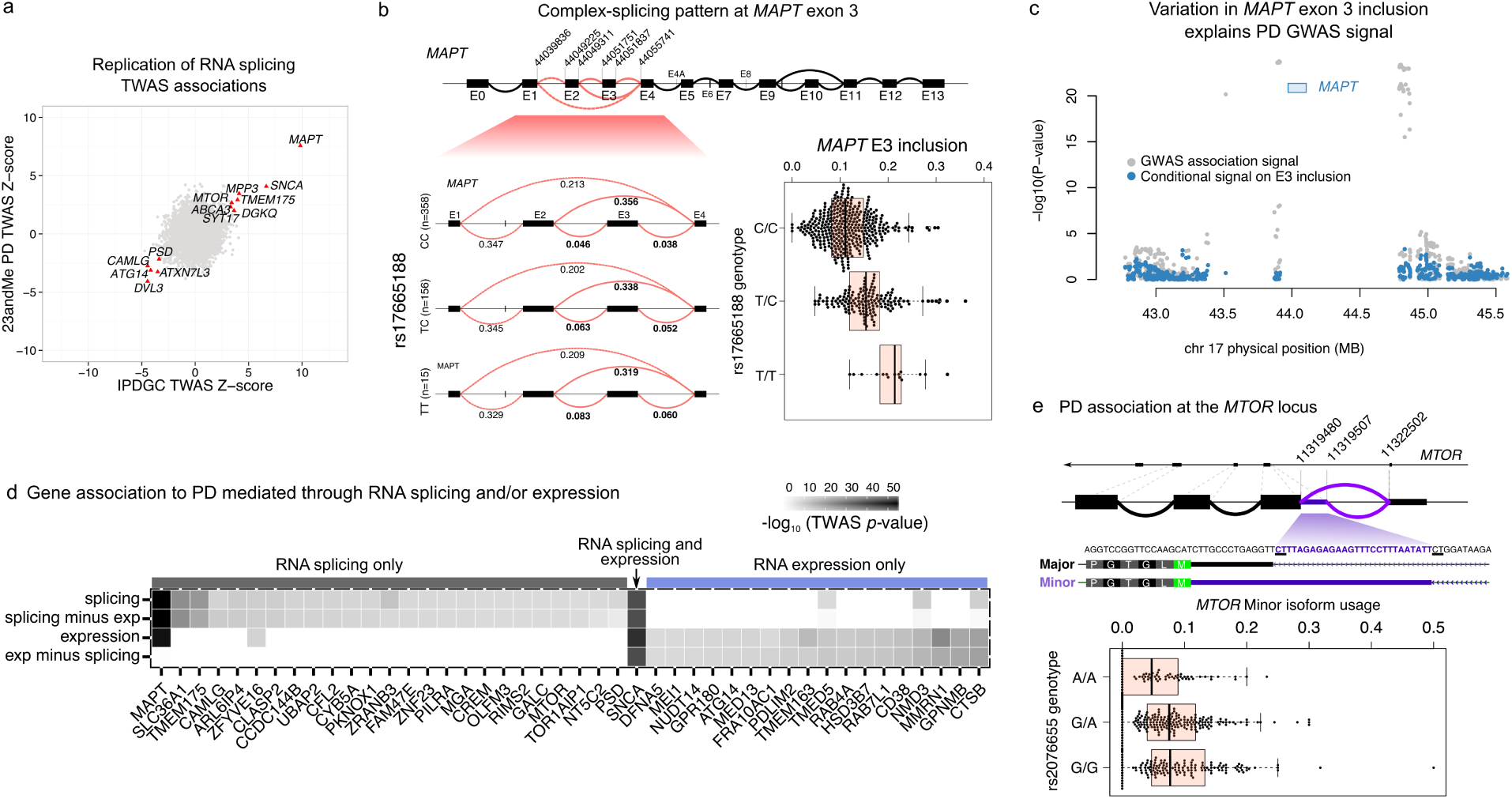
PD genetic variants affect splicing of nearby genes. **(a)** Replication of TWAS Z-score across two PD cohorts. **(b)** The H1/H2 tag SNP rs17665188 is associated with MAPT exon 3 inclusion. The T allele, which tags the H2 haplotype, doubles inclusion of exon 3 and is associated with increase risk for PD. **(c)** PD TWAS signal at the MAPT locus (grey) and TWAS signal after removing effect of MAPT exon 3 inclusion (cyan). This analysis shows that the association is largely explained by MAPT exon 3 inclusion. (d) Heatmap of genes identified from out TWAS analysis using imputed RNA splicing and expression (splicing and expression respectively). The rows “splicing minus exp” and “exp minus splicing” denote association strength after conditioning on expression and splicing, respectively. *SNCA* is associated to PD through both a genetic effect on RNA splicing and expression. **(d)** PD-associated variants at MTOR locus are associated to an increase in a minor MTOR isoform that extends the 5’ of an exon in the 5’ UTR.

### PD genetic variants often affect splicing of nearby genes

To obtain a better understanding of the associations identified in our TWAS, we next focused on dissecting the mechanisms by which specific genes were associated to PD. We identified many known PD genes, such as *MAPT*, the microtubule-associated protein *tau* (chr17:44049311:44055741; *p* = 2.60 × 10^−24^, Fig. 4b) and *SNCA* (chr4:90756843:90757894; *p* = 1.70 × 10^−16^) that were associated to PD through RNA splicing. However, several genes, including *MAPT*, were found in both our RNA expression and splicing TWAS analyses. This finding is consistent with the existence of contradictory studies that either propose that differences in *MAPT* expression^11^, or that differences in *MAPT* splicing (exon 3 inclusion^40^ or exon 10 inclusion^11^) is driving the PD genetic association at this locus. We therefore asked whether the associations we identified were more likely to be mediated through RNA splicing or through RNA expression, and whether we could predict which gene or splicing event were most likely to be causal. At the *MAPT* locus, six associations to splicing events in 3 genes were identified. Using FUSION^25^, we found that only one splicing event (inclusion of *MAPT* exon 3) remained significant (conditional P-value < 2.60 × 10^−24^; Fig. 4c,d; Supplementary Table 3) after conditioning on all other association signals, including *MAPT* total RNA expression levels. To identify which genetic variant is associated to *MAPT* exon 3 inclusion, we searched our splicing QTL data from DLPFC and found that a nearby SNP (rs17665188/chr17:44357351) was strongly associated to exon 3 inclusion levels. Importantly, this SNP tags (*r* > 0.93) two well-known haplotypes (H1/H2)^40^. Haplotype H2 is associated with increased *MAPT* exon 3 inclusion in DLPFC (*p* < 2.2 × 10^−16^, LR *t*-test). Therefore, we conclude that splicing variation of *MAPT* exon 3 in the brain likely explains the reported association between the H1/H2 haplotypes and PD^41^.

While some of the genes we identified were located within loci previously identified using GWAS, we discovered 47 associations that were in novel PD loci (Supplementary Table 1). Most of these were located in loci with suggestive associations in PD GWAS (5 × 10^−8^ < *p* < 1 × 10^−6^, Supplementary Figs. 4-42), and four genes (*CTSB*, *PDLIM2*, *GALC*, and *C8orf5*) were found to be genome-wide significant in the most recent PD GWAS meta-analysis ^3^. One of these genes is cathepsin B (*CTSB*), which is a part of protease essential in *α*-synuclein lysosomal degradation^42^. We also detected *MTOR* as a novel candidate PD gene (Supplementary Fig. 5). *MTOR* is a highly conserved serine/threonine kinase expressed in most of mammalian cell types, and plays a central role in regulation of autophagy^43,44^. Of note, recent data have shown that dysregulation of mTOR is implicated in the pathogenesis of PD^45,46^, and it has been suggested as a novel therapeutic target for PD. Here, we found that a putative PD risk SNP (rs207655) was associated to a significant increase in usage of a rare *MTOR* isoform (Fig. 4e). Overall, our PD TWAS in DLFPC highlights the convergence of old and new PD genes in autophagy/lysosomal degradation pathways.

### Enrichments of PD TWAS genes in functional pathways

Lastly, we hypothesized that the newly identified TWAS genes may be part of the same network or pathway as known PD susceptibility genes. To test this, we used GeNets^47^ to measure the protein-protein interaction (PPI) network connectivity of our TWAS genes with known PD susceptibility genes. First, as one might expect, we found that monogenic PD genes form a PPI network that is directly connected (i.e., they form shared communities) to genes in PD GWAS loci (*p* < 7.2 × 10^−3^) (Fig. 5a). Remarkably however, when we incorporated the TWAS genes with known PD susceptibility genes (including the monogenic genes), we found an expanded PPI network with four distinct communities of genes encoding for proteins that physically interact (Fig. 5b; *p* < 2.3 × 10^−3^). The genes in the TWAS-prioritized PPI network are highly enriched for biological pathways including lysosome (Bonferroni *p* = 0.0017) and α-synuclein aggregation pathways (Bonferroni *p* = 1.8 × 10^−5^) (Fig. 5b). These results support our hypothesis that the novel candidate PD genes we identified in this study are part of a larger set of interacting genes with coherent biological function, of which the lysosome pathway may be the most central in PD.

**Figure 5:**
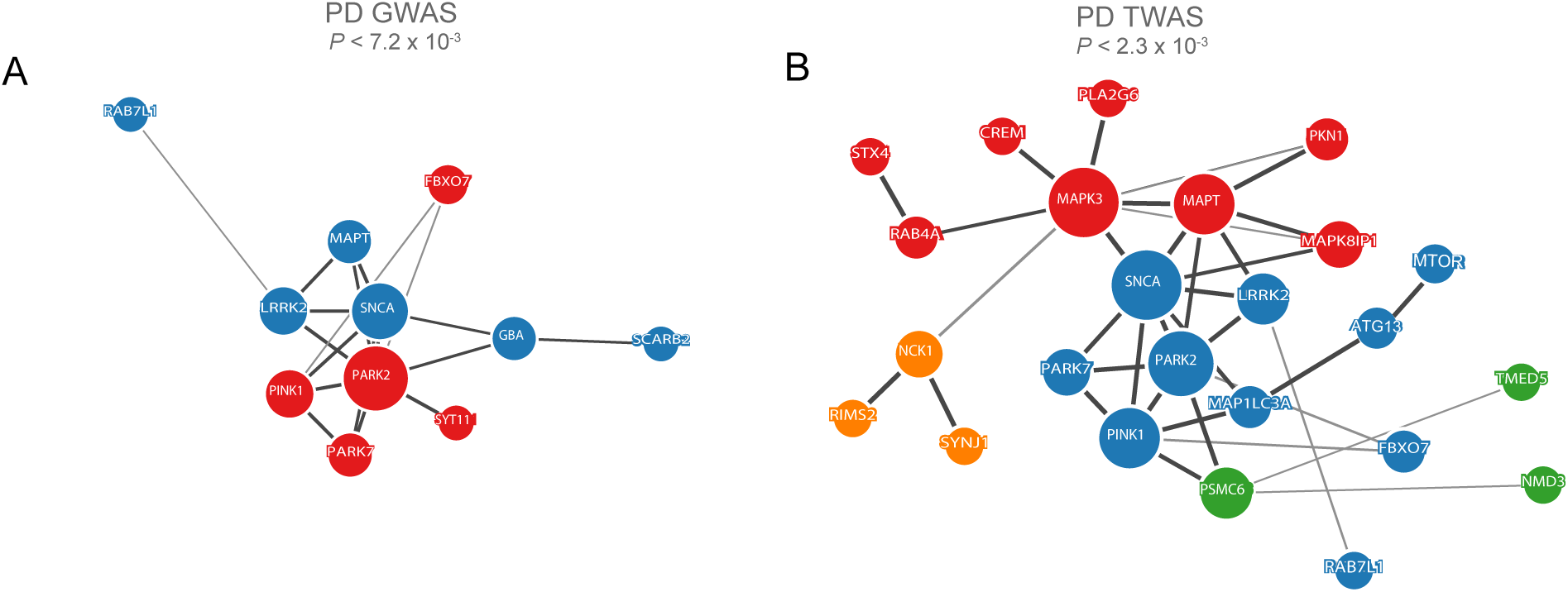
Proteins nominated by PD TWAS form expanded PPI and are enriched in lysosomal pathway. (**a**) Proteins product of known PD GWAS and monogenic genes form significant (*p* < 7.2 × 10^−3^) PPI network with two communities. (**b**) Protein product nominated by PD TWAS and monogenic genes form a significantly expanded PPI (*p* < 2.3 × 10^−3^) network with known PD genes with four communities.

## Discussion

While a growing body of evidence has implicated innate immunity mechanisms in PD pathophysiology, it remains unclear which genes and specific pathways are involved. Our analysis of gene expression in peripheral monocytes suggests that PD-associated genetic risk factors influence innate-immune mechanisms. However, more work is required to determine whether we observed these associations in peripheral monocytes because they are contributing to PD directly via infiltrating cells that differentiate into macrophages, or because these associations reflect resident microglial activity in the brain (peripheral cells may serve as a related, proxy myeloid cell type). Further, we leveraged PD GWAS data and large-scale transcriptomic data from human cortex to identify genes for which the genetic component of expression level or differential splicing is associated with PD. Our transcriptome-wide association study corroborated many of the known PD genes, but also identified several candidate disease genes in novel PD loci. Our observations imply gene expression and splicing regulation in neuronal tissue as key mechanisms that mediate genetic risk for PD. Novel genes (e.g., *CTSB* and *MTOR*) identified in our study increase our understanding of the pathways that drive PD biology and further highlight the importance of lysosomal function in the etiology of PD.

Overall, our study prioritizes genes for subsequent experimental follow-up, which will help interrogate the molecular mechanisms underlying PD. Moreover, our catalog of splicing QTLs in DLFPC are made available with this study (see URLs), which provides a starting point for further mechanistic work to elucidate the role of associated genes in PD. Thus, this work represents a significant step towards understanding the genetic basis of PD.

## Online Methods

### Data acquisition, quality control, and normalization

#### Transcriptomics Atlases for LDSC-SEG

Data for LDSC-SEG was prepared as described in^19^. The following data were used in our LDSC-SEG analyses: (1) GTEx samples that had more than 4 samples with at least one read count per million and at least 100 genes with at least one read count per million were TPM normalized^48^. (2) Publicly available data for DEPICT^14^ was downloaded and pruned so that no two tissues had *r*^2^ > 0.99. Two sets of highly correlated tissues (eyelid, conjunctiva, anterior eye segment, tarsal bones, foot bones, and bones of the lower extremity; and connective tissue, bone and bones, skeleton, and bone marrow) were removed completely, leaving 152 tissues. (3) Affymetrix GeneChip expression array data from mouse forebrain sorted cells was downloaded from GEO (GSE9566)^20^. (4) Publicly available gene expression data from the ImmGen project^15^ was downloaded from GEO (GSE15907, GSE37448). Ensembl orthologs were used to map to human genes.

#### Transcriptomics Panels for TWAS

The following panels were used in our study: (1) Common Mind consortium (CMC) RNA-seq data generation and processing were previously described in^18^. Briefly, dorsolateral prefrontal cortex (Brodmann areas 9/46) was dissected from post-mortem brains of 258 individuals with schizophrenia and 279 control subjects. These individuals were of diverse ancestry, had no Alzheimer’s Disease or Parkinson’s Disease neuropathology, had no acute neurological insults (anoxia, stroke, or traumatic brain injury) immediately before death, and were not on ventilators near the time of death. Total RNA was isolated from homogenized tissue and rRNA depleted. 100bp paired-end reads were obtained using an Illumina HiSeq 2500, and mapped using TopHat. Genotyping was performed using the Illumina Infinium HumanOmniExpressExome-8 v 1.1b chip. (2) Fairfax et al. monocyte expression data generation was previously described in^16^. In short, blood was collected from 432 individuals of European ancestry, and CD14^+^ monocytes were isolated from PBMCs via MACS. RNA was quantified using the Illumina HumanHT-12 v4 BeadChip. Expression was normalized and transformed using robust spline normalization in R using the Lumi package, and corrected for batch effects using the ComBat package. Using a linear model for the effect of incubation time on expression, incubation time was regressed out of the normalized expression values. Genotyping was performed using the Illumina HumanOmniExpress-12 v1.0 chip. (3) Cardiogenic monocyte expression data generation was previously described in^49,17^. Monocytes were sorted from whole blood from individuals of European descent, and expression was assessed from RNA using the Illumina HumanRef 8 v3 Beadchip. Individuals were genotyped on Illumina Human 610 Quad custom arrays. The gene expression data is available via https://ega-archive.org/studies/EGAS00001000411. (4) ROS/MAP RNA-seq data generation was previously described in^22,21^. ROS/MAP is a prospective cohort of aging individuals, where individuals are healthy at enrollment and 38% have clinical Alzheimer’s disease at the time of death. Dorsolateral prefrontal cortex grey matter was dissected from 540 post-mortem brains. Sequencing libraries were prepared using a strand-specific dUTP protocol with poly-A selection, and sequencing on an Illumina HiSeq produced 101bp paired-end reads. Individuals were genotyped on an Affymetrix GeneChip 6.0.

#### GWAS Datasets

We performed transcriptome-wide association study using PD GWAS summary statistics from Nalls et al.^27^. For discovery, we restricted our GWAS from PD cases and controls to IPDGC, PD GWAS Consortium, CHARGE, PDGENE, and Ashkenazi studies cohorts (9581 cases and 33245 controls). For replication, we used the summary statistics from 23andMe subsets only (v2 and v3) from Nalls et al.^27^ (4124 cases and 62037 controls).

#### Transcriptome-wide Association Studies

Transcriptome-wide Association Studies (TWAS) is a powerful strategy that integrates SNP-expression correlation (cis-SNP effect sizes), GWAS summary statistics and LD reference panels to assess the association between cis-genetic component of expression and GWAS ^25^. TWAS can leverage large-scale RNA-sequencing data to ?impute? tissue-specific genetic expression levels from genotypes (or summary statistics) in larger samples, which can be tested to identify potentially novel associated genes ^26^,^25^.

We used RNA-seq data and genotypes from CommonMind Consortium to impute the cis genetic component of expression/intron usage into large-scale late-onset PD GWAS of IPDGC as discovery. For replication 23andMe The complete TWAS pipeline is implemented in FUSION (http://gusevlab.org/projects/fusion/) suite of tools. The details steps implemented in FUSION are: (1) estimate heritability of gene expression or intron usage unit and stop if not significant. We estimated using a robust version of GCTA-GREML^50^, which generates heritability estimates per feature as well as the as well as the likelihood ratio test (LRT) P-value. Only features that have heritability of Bonferroni-corrected P < 0.05 were retained for TWAS analysis. (2) The expression or intron usage weights were computed by modeling all cis-SNPs (1MB +/− from TSS) using best linear unbiased prediction (BLUP), or modeling SNPs and effect sizes (BSLMM), LASSO, Elastic Net and top SNPs. A cross-validation for each of the desired models are performed; (3) Perform a final estimate of weights for each of the desired models and store results. The imputed unit is treated as a linear model of genotypes with weights based on correlation between SNPs and expression in the training data while accounting for LD among SNPs. To account for multiple hypotheses, we applied a false discovery rate of 5% within each expression and splicing reference panel that was used.

#### Joint and conditional analysis

Joint and conditional analysis was performed using the summary statistic-based method described in ^51^, which we applied to genes instead of SNPs. We used TWAS association statistics from the main TWAS results and a correlation matrix to evaluate the joint/conditional model. The correlation matrix was estimated by predicting the cis-genetic component of expression for each TWAS gene and computing Pearson correlations across all pairs of genes. We used FUSION tool to perform the joint/conditional analysis, generate conditional outputs, and generate plots.

#### Splicing QTL Mapping

We used Leafcutter^38^ to obtain the proportion of intron defining reads to the total number of reads from the intron cluster it belongs to. This intron ratio describes how often an intron is used relative to other introns in the same cluster. We used WASP ^52^ to remove read-mapping biases caused by allele-specific reads. This is particularly significant when a variant is covered by reads that also span intron junctions as it can lead to a spurious association between the variant and intron excision level estimates. We standardized the intron ratio values across individuals for each intron and quantile normalize across introns^24^ and used this as our phenotype matrix. We used linear regression (as implemented in fastQTL^53^) to test for associations between SNP dosages (MAF 0.01) within 100kb of intron clusters and the rows of our phenotype matrix that correspond to the intron ratio within each cluster. As covariate, we used the first 3 principal components of the genotype matrix to account for the effect of ancestry plus the first 15 principal components of the phenotype matrix (PSI) to regress out the effect of known and hidden factors. To estimate the number of sQTLs at any given false discovery rate (FDR), we ran an adaptive permutation scheme ^53^, which maintains a reasonable computational load by tailoring the number of permutations to the significance of the association. We computed the empirical gene-level p-value for the most significant QTL for each gene. Finally, we applied Benjamini-Hochberg correction on the permutation p-values to extract all significant splicing QTL pairs with an FDR < 0.05.

#### Colocalization

We used coloc 2.3-1^39^ to colocalize the PD association signal at TWAS loci with QTL signals. For each locus, we examined all SNPs available in both datasets within 500 Mb of the SNP identified in TWAS as the top QTL SNP, and ran coloc.abf with default parameters and priors. We called the signals colocalized when (coloc H3 + H4 ≥ 0.8 and H4/H3 ≥ 2).

#### GWAS Enrichment

We used GARFIELD to test for enrichment of IGAP AD GWAS SNPs among splicing QTLs and other publicly available QTL datasets^54^. GARFIELD performs greedy pruning of GWAS SNPs (LD *r*^2^ >0.1) and then annotates them based on functional information overlap. It quantifies fold enrichment at GWAS *P* < 10^−5^ significant cutoff and assesses them by permutation testing, while matching for minor allele frequency, distance to nearest transcription start site and number of LD proxies (*r*^2^ > 0.8) ^55^.

#### Protein-protein Interaction Network and Pathway Analysis

We constructed a protein-protein interaction (PPI) network using the online tool GeNets (https://apps.broadinstitute.org/genets) to determine whether the PD TWAS genes significantly interact with each other and with known (Mendelian genes) PD associated proteins. GeNets creates networks of connected proteins using evidence of physical interaction from the InWeb database, which contains 420,000 high-confidence pair-wise interactions involving 12,793 proteins^47^. Community structures of the underlying genes are displayed in GeNets. Communities are also known as modules or clusters. Community feature highlights genes that are more connected to one another than they are to other genes in other modules. To assess the statistical significance of PPI networks, GeNets applies a within-degree node-label permutation strategy to build random networks that mimic the structure of the original network and evaluates network connectivity parameters on these random networks to generate empirical distributions for comparison to the original network. In addition to PPI network analysis, GeNets allows for gene set enrichment analysis on genes within the PPI network. We used Molecular Signatures Database (MSigDB) Curated Gene Sets (C2), curated from various sources such as online pathway databases, the biomedical literature, and knowledge of domain experts and Canonical Pathways (CP), curated from pathway databases such KEGG, BioCarta, Reactome, etc. to test for gene set enrichment within the PPI network. Then a hypergeometric testing is applied to get P-value for gene set enrichment. We used Bonferroni-corrected *P* < 0.05 to correct for multiple hypothesis testing.

## Acknowledgement

This work was supported by the US National Institutes of Health (NIH grant R01AG054005). This work was supported in part through the computational resources and staff expertise provided by Scientific Computing at the Icahn School of Medicine at Mount Sinai. We would like to thank Elisa Navarro, Satesh Ramdhani, Hae Kyung Im, and Tim Ahfeldt for their insightful comments on the paper. We thank the patients and families who donated material for CommonMind Consortium data. The CommonMind Consortium data are available in CMC Knowledge Portal: https://www.synapse.org/#!Synapse:syn4923029. Data were generated as part of the CMC supported by funding from Takeda Pharmaceuticals Company Limited, F. Hoffman-La Roche Ltd and NIH grants R01MH085542, R01MH093725, P50MH066392, P50MH080405, R01MH097276, RO1-MH-075916, P50M096891, P50MH084053S1, R37MH057881 and R37MH057881S1, HHSN271201300031C, AG02219, AG05138 and MH06692. Brain tissue for the study was obtained from the following brain bank collections: the Mount Sinai NIH Brain and Tissue Repository, the University of Pennsylvania Alzheimer’s Disease Core Center, the University of Pittsburgh NeuroBioBank and Brain and Tissue Repositories and the NIMH Human Brain Collection Core. CMC Leadership: Pamela Sklar, Joseph Buxbaum (Icahn School of Medicine at Mount Sinai), Bernie Devlin, David Lewis (University of Pittsburgh), Raquel Gur, Chang-Gyu Hahn (University of Pennsylvania), Keisuke Hirai, Hiroyoshi Toyoshiba (Takeda Pharmaceuticals Company Limited), Enrico Domenici, Laurent Essioux (F. Hoffman-La Roche Ltd), Lara Mangravite, Mette Peters (Sage Bionetworks), Thomas Lehner, Barbara Lipska (NIMH).

We thank the participants of ROS and MAP for their essential contributions and gift to these projects. The ROS/MAP data are available at the The Rush Alzheimer’s Disease Center (RADC) Research Resource Sharing Hub at www.radc.rush.edu. The ROS/MAP and MSBB mapped RNA-seq data that support the findings of this study are available in AMP-AD Knowledge Portal (https://www.synapse.org/#!Synapse:syn2580853) upon authentication by the Consortium. This work has been supported by many different NIH grants: P30AG10161, U01AG046152, R01AG16042, R01AG036836, R01AG015819, R01AG017917, R01AG036547.

## Author Contributions

T.R. conceived of the project. Y.I.L., G.W., and T.R. performed the analyses. T.R. and Y.I.L. wrote the manuscript.

## URLs

DLPFC sQTL browser: https://leafcutter.shinyapps.io/sQTLviz/.

